# FindBacksplice: a Tool for Locating Circular RNA Backsplice Coordinates

**DOI:** 10.1101/2025.09.08.674962

**Authors:** Matthew Kraljevic, Arzu Ozturk, Marietta Jank, Richard Keijzer, Richard LeDuc

**Affiliations:** Department of Department of Biochemistry and Medical Genetics, University of Manitoba and Children’s Hospital Research Institute of Manitoba, Winnipeg, MB, Canada; Divison of Pediatric Surgery, Department of Surgery, Max Rady College of Medicine, University of Manitoba, and Children’s Hospital Research Institute of Manitoba, Winnipeg, MB, R3A 1S1, Canada; Department of Pediatric Surgery, University Medical Center Mannheim, Heidelberg University, Mannheim, Germany

## Abstract

Circular RNAs (circRNAs) are generated through back-splicing, a process where a backsplice junction (BSJ) is formed, based on the circRNAs’ unique sequence. BSJs are highly conserved and can be mapped to chromosomal coordinates. Current platforms determine these coordinates from bulk RNA sequencing data. We aimed to develop a tool capable of determining backsplice coordinates based on BSJ sequences for any species and genome version.

**Motivation:** circRNAs are emerging as an important regulator of cellular differentiation and other biologically important processes. Common tools for circRNA analyses require a circular RNA’s backsplice coordinates. These can be accessed from public databases. However, the coordinates are specific to a version of a species’ genome, and are unavailable for many model organisms.

**Results:** We have developed a Python-based, command line tool, FindBacksplice, which produces backsplice coordinates for any available genome, based on a circRNA’s BSJ sequence. Implemented in Python, this script is integrated with BLAST for use in existing pipelines. We were able to find valid locations of backsplices for known human BSJs in the rat genome and produce backsplice coordinates for use in existing pipelines.

**Availability and implementation:** FindBacksplice is available at github.com/m-kraljevic/findbacksplice

## 1 Introduction

Known from the 1970s, circular RNAs (circRNA) are stable, long, non-coding RNAs characterized by a loop structure and emerging as potential biomarkers of human disease (Sanger et al. 1976, Patop et al. 2019, Liu & Chen 2022). CircRNA molecules are formed via a variety of molecular pathways across the tree-of-life, but those created by backsplicing mRNA are of particular interest in translational research. These circRNAs are composed of a single-stranded loop of RNA, which is enzymatically formed with a covalent bond between a downstream splice donor site and an upstream acceptor site. This formation event is referred to as “back-splicing” (Kristensen et al., 2019). The donor and acceptor sites are highly evolutionarily conserved allowing candidate circRNA molecules to be inferred from within a sequenced genome based on the backsplice junction sequence (BSJ) (Varela-Martínez et al. 2021, Chuang et al. 2023, Zou et al. 2025).

Over 100,000 circular RNAs are known. Initially considered a result of abnormal RNA splicing, circular RNAs have since been shown to play a role in governing cellular processes and disease pathogenesis. Given the evolutionarily conserved nature of backslices, most human backsplice-derived circRNAs are also present in model organisms (Kristensen et al., 2019).

Tools like CRAFT, circRNAProfiler, CiLiQuant, circNetVis, circR, and circMiMi require the BSJ coordinates of circular RNAs as input (Aufiero et al. 2020, Chiang et al. 2022, Dal Molin et al. 2022, Dori et al. 2022, Morlion et al. 2022, Nguyen et al. 2024): these are the coordinates of the donor and acceptor sequences within the genome. These coordinates are also required for the development of PCR primers to detect the transcription of exonic circular RNAs within experimental samples (Chuang et al., 2023). BSJ coordinates for a specific circRNA within a given genome may be retrieved from databases such as circBase (Glažar et al., 2014) or circAtlas (Wu et al., 2023) or may be provided by circRNA microarray manufacturers (e.g. Arraystar Inc.). However, these coordinates are always associated with a single genome version of the organism and the given combination of backsplice sequences.

Several tools such as circExplorer2 and Find_circ, (Memczak et al. 2013, Ma et al. 2021) empirically locate backsplices from bulk RNA data. Typically, existing tools take bulk RNA seq data and use the unmapped high-quality reads to determine if any are consistent with spanning a known backsplice. If they are, these tools then return the coordinates for that backsplice.

We present a novel bioinformatic tool, FindBacksplice, which determines backsplice coordinates for a user-supplied genome based on a BSJ input sequence. This addresses the reasonably simple but important task of finding the backsplice genomic locations for the BSJ sequence, so that for a given sequence, backsplice coordinates may be produced for use with other analysis tools. Our algorithm enables locating backsplices between species in any user supplied genome. Additionally, backsplice locations provided by a database or PCR supplier may not be based on a genome version used in a researcher’s pipeline. FindBacksplice ensures backsplice coordinates are appropriate for any genome.

FindBacksplice allows BSJ primers to be translated into genomic locations in the genome of any species, allowing tools to be applied to model organisms and filling a gap in developing pipelines for circRNA analysis. This tool enabled the analysis of a rat model of congenital diaphragmatic hernia (Jank et al., 2024). The coordinates determined by this tool can serve as input for any platform that investigates the biological significance of circRNA functions. For example, tools such as CRAFT and CircMiMi infer biological processes based on predicted circRNA interactions with RNA binding proteins, microRNAs and messenger RNAs using ontological analyses and gene set enrichment analysis.

FindBacksplice functions as a Python-based command line solution to be integrated into other existing CircRNA pipelines, and the code is freely available at https://github.com/m-kraljevic/FindBacksplice.

## 2 Tool Description

FindBacksplice outputs a.csv file containing valid chromosomal backsplice locations in a user-specified organism for circRNAs determined from their BSJ sequence. It is a command line tool written in Python. It was designed to be integrated with the BLAST+ (Camacho et al., 2009) implementation of BLAST (Altschul et al., 1990) and it is recommended to be run with the output of command-line-based nucleotide BLAST (2021), but may also use the XML output of BLAST web portals. Being run from a terminal and utilizing commonplace software such as Python and BLAST, FindBacksplice is suitable for use in circRNA analysis pipelines which use backsplice coordinates as input (such as CRAFT (Dal Molin et al. 2022), circMiMi (Chiang et al. 2022) and ciLiQuant (Morlion et al. 2022).

## 3 Real usage case and algorithm explained

FindBacksplice implements two methods to produce backsplice coordinates. The first exclusively utilizes the output of BLAST. The second uses a combination of BLAST output and BioPython (Cock et al., 2009) to parse the reference genome.

The BLAST-only approach is seen when searching for the backsplice coordinates for hsa_circRNA_406614 in rat genome version 7.2, Figure 1. Searching the query (Figure 1A) with BLAST against the reference genome yields two locations (Figure 1B) which together comprise the entire BSJ sequence. The start of the BSJ sequence is downstream from the end of the BSJ sequence which implies that the returned location represents a valid BSJ in the genome (Figure 1C). For this example, FindBacksplice will return chr2:35536665-35561195.

**Figure 1.**
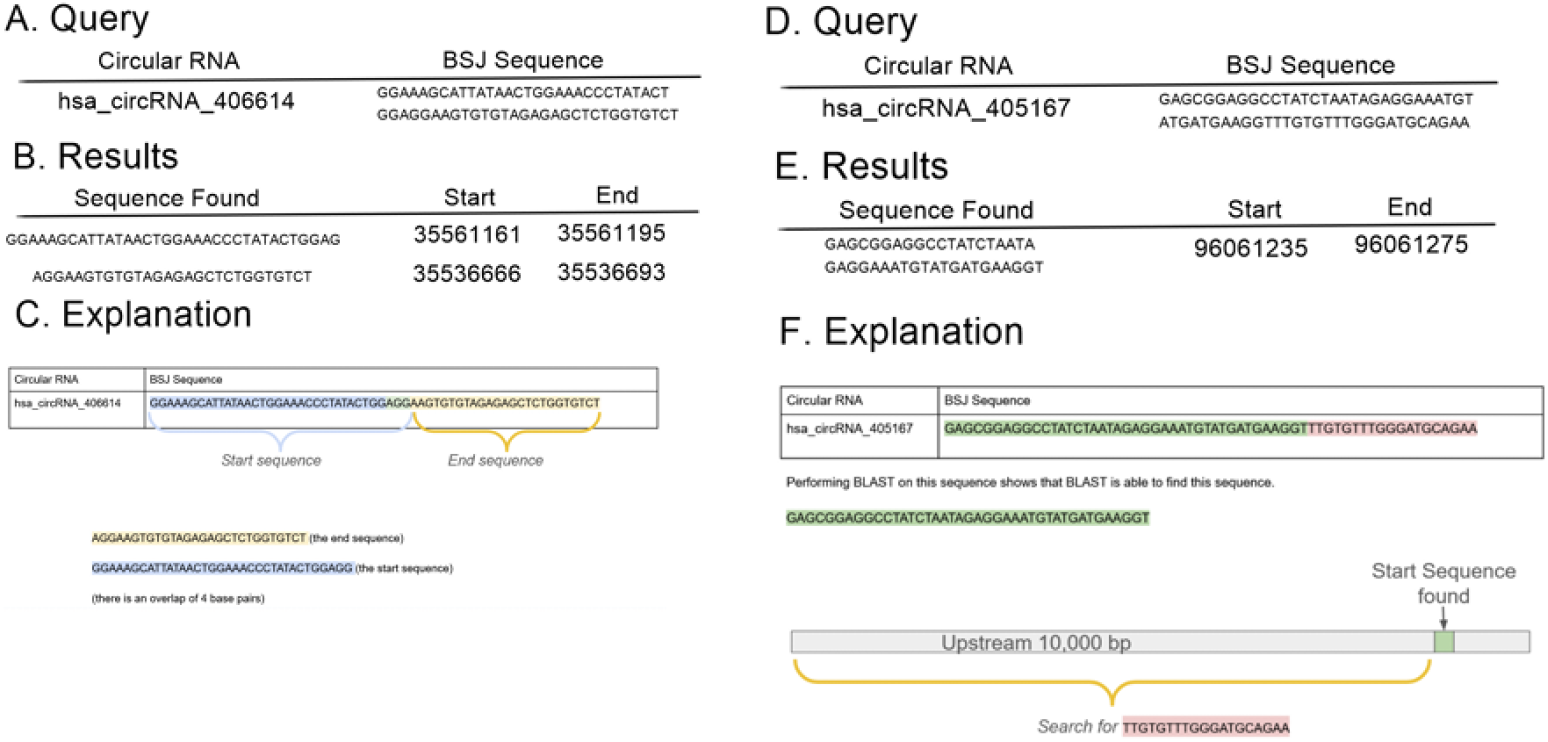
An ideal and problematic example for determining backsplice coordinates for human circRNA in the rat genome. A) The human circRNA sequence for an idyllic example of using BLAST. B) The two results returned, and C) a visual representation of the two halves of the backsplice with the start at the end, and the end at the start. D) The human sequence for a problematic example of using BLAST. E) Only one result is returned, and F) a visual representation of the need to search for the remaining 19 bp upstream of the located sequence.5055

Unfortunately, the donor or acceptor sequences may be too short for BLAST to locate. This is seen when searching hsa_circRNA_405167 in rat v7.2, Figure 1D. Performing BLAST on this sequence returns a partial sequence of the BSJ.

In this case, only the first 41 nucleotides of the BSJ sequence (Figure 1E) were returned, the remaining 19 basepairs (bp) were missed. BLAST is most effective when searching for sequences longer than 22 bp long (EBI Web Services). Due to this limitation, we implement an alternate solution.

This solution uses BioPython to load a user-defined number of basepairs related to the sequence returned by BLAST, Figure 1F. In this example, 96,061,235 to 96,051,235 on chromosome 15 on the reference genome. The default search space length is 10,000 as circRNA molecules are typically under 1,000 bp in length (Ding et al, 2018), but this is left user-defined, as there exists evidence of circRNAs reaching 81,000 bp in length (Margvelani et al., 2024). Using the Python index function, the tool attempts to locate the remaining 19 bp (TTGTGTTTGGGATGCAGAA) of the BSJ sequence.

The remaining 12 bp sequence of hsa_circRNA_405167 are found at location 96,058,577 – 96,058,596 in rat. This yields chr15:96058577-96061275 as the candidate rat location for the backsplice of hsa_circRNA_405167.

These examples are all based on positive strand sequences. If BLAST returns sequences on the negative strand, the same algorithm is applied, but FindBacksplice will search the genome in the opposite direction for the reverse complement of the sequence. The final output will show the strand as negative for these backsplice coordinates.

## 4 System Requirements

FindBacksplice is compatible with Windows, macOSX, and Linux.

FindBacksplice requires Python 3.0 or greater and the packages BioPython, argparse and pandas. Package versions used in the development of FindBacksplice were BioPython v1.85 (Cock et al., 2009), argparse v1.1 (Python Software Foundation) and pandas v2.3.2 (McKinney, 2010).

The installation of these libraries may vary depending on the user’s Python installation. Users can refer to our Github repository for installation troubleshooting for different Python environments.

### Preparing Input

FindBacksplice requires three files to run:

i. File containing all BSJ Sequences in.fasta format
ii. Genome file in.fasta format
iii. BLAST output in.xml format

FindBacksplice was developed using command line BLAST 2.16.0+ (Camacho et al., 2009) on a Red Hat Enterprise Linux environment. The BLAST database for each species was created as:

**makeblastdb -in species_genome.fasta dbtype nucl -out db/species**

The XML output was generated as:

**blastn -db db/species_db -query bsj_sequences.fasta -outfmt 5 > blast_output.xml**

NOTE: web portal implementations of BLAST (National Center for Biotechnology Information) can provide output in XML format, but this output must be saved locally as an.xml file to be available to FindBacksplice.

## 5 Installation and Use

i. Visit the GitHub repository for FindBacksplice, https://github.com/m-kraljevic/FindBacksplice, and download or clone the repository. If downloaded, unzip the FindBacksplice.zip file.
ii. From command line, navigate to the directory containing FindBacksplice.py
iii. Run the script with the required input files. For example

python FindBacksplice.py -i /path/to/bsj_sequences.fasta -g /path/to/genome.fasta -b /path/to/blast_output.xml -o /path/to/backsplice_coordinates.csv

The -o flag is used to denote the location and name of the output csv file, which will default to the same folder as FindBacksplice.py.

A help flag is available by running python FindBacksplice.py -h to provide more detailed information on the flags and input parameters.

## 6 Example use case and output

Here we use FindBacksplice to locate a backsplice conserved between rat and human. A sample file, conserved_sequence.fasta was created containing one BSJ sequence.

> hsa_circRNA_006314

ACAGGATATGCCTTGTCAGATCTGCTACTTGAACTACCCTAACTCGAATCCAGCAACTAT

For demonstration purposes, FindBacksplice.py file was run twice, first on human and then on rat.

python FindBacksplice.py -i conserved_sequence.fasta -g Homo_sapiens.GRCh38.dna.primary_assembly.fa -b human_blast_output.xml -o human_backsplice_coordinates.csv

python FindBacksplice.py -i conserved_sequence.fasta -g Rattus_norvegicus.mRatBN7.2.dna.toplevel.fa -b rat_blast_output.xml -o rat_backsplice_coordinates.csv

These produced two.csv files containing the following information

human_backsplice_coordinates.csv

chr5,65242094,65273449,-

rat_backsplice_coordinates.csv

chr2,35536666,35561195,+

### Blast database generation

To generate the BLAST input files needed for the above example, the input BSJ sequence file was run against two blast databases produced with:

Homo_sapiens.GRCh38.dna.primary_assembly.fa downloaded from Ensembl on July 24, 2024 (Harrison et al., 2023)

Rattus_norvegicus.mRatBN7.2.dna.toplevel.fa downloaded from Ensembl on October 30, 2024 (Harrison et al., 2023)

The databases were created using the commands

makeblastdb -in Homo_sapiens.GRCh38.dna.primary_assembly.fa dbtype nucl -out db/human

makeblastdb -in Rattus_norvegicus.mRatBN7.2.dna.toplevel.fa dbtype nucl -out db/rat

### Blast XML production

The blast output was produced with the commands

blastn -db db/human -query conserved_sequence.fasta -outfmt 5 > human_blast_output.xml

blastn -db db/rat -query conserved_sequence.fasta -outfmt 5 > rat_blast_output.xml

## 7 Validation

To validate FindBacksplice, a test dataset of backsplice sequences was generated using the circRNADb database of known human circRNA coordinates (based on Human Genome hg19) and circRNA sequences (Chen et al., 2016). The 32,914 circRNAs available were filtered to only those with coordinates spanning under 10,000 bp and greater than 60 bp to represent the default parameters of the tool. Then, 1000 of these filtered circRNAs were chosen at random.

To generate BLASTable BSJ sequences, a theoretical BSJ was generated for each of the 1000 randomly chosen circRNAs. This was done by concatenating the last 30 bp of the circRNA sequence to the first 30 bp of the circRNA sequence.

BLAST was then run on these 1000 theoretical BSJ sequences using human genome hg19 accessed from Ensembl (Harrison et al., 2023) and the blast output, genome and sequences were subsequently inputted into FindBacksplice through command line with default settings. Of the 1000 inputted sequences, 951 backsplice coordinates were generated on the correct chromosome and strand within 22 bp of the coordinates reported by circRNAdb. 7 were reported with different chromosomes and location, and 42 were unable to be found by the tool.

## 8 Conclusions

FindBackSplice is a useful tool for bioinformatic workflows exploring the role of circRNAs in model organisms. It handles several special cases that are found when mapping BSJ sequences to a new genome including when one or the other end of a backsplice is too short to be located with BLAST and when the splice junction occurs on the negative strand. This tool enables the use of many other bioinformatic tools written for researching circRNAs in humans, to now be applied to model organisms. The tool is written so it can be easily used by an analyst performing a “one-off” analysis or it can be incorporated into bioinformatic pipelines.

## Supplementary data

All code and validation data is available in our Github repository: github.com/m-kraljevic/findbacksplice

